# Continuous pharyngeal endoderm links external and internal gills

**DOI:** 10.64898/2026.08.12.744359

**Authors:** Himanshi Singh, Michaela Kavková, Jan Vintr, Lorena Agostini Maia, Jakub Harnoš, Jan Krivanek, Radek Šindelka, Vladimír Soukup

**Author notes:** Author of correspondence: Vladimír Soukup.

## Abstract

Amphibians develop both external and internal gills during ontogeny, offering an opportunity to investigate the developmental relationship between these positionally distinct respiratory organs. Although internal gills of vertebrates are widely accepted to arise from pharyngeal endoderm, external gills have long been regarded as purely ectodermal outgrowths, obscuring their relationship to other vertebrate gills. Here, we combine histological analysis with direct lineage tracing in the Mexican axolotl (*Ambystoma mexicanum*) and the African clawed frog (*Xenopus laevis*) to resolve the embryonic origin of amphibian gills. We show that the external gill develops as a continuous epithelial extension of the pharyngeal endoderm, which forms its basal epithelium and reaches the distal gill tip. In the frog, this extension remains continuous with the epithelium giving rise to the internal gills. Rather than representing separate epithelial structures, external and internal gills therefore arise from a shared epithelial domain of the pharyngeal endoderm. These findings resolve a longstanding question concerning the embryonic origin of amphibian gills and provide a developmental viewpoint for understanding how spatially diverse vertebrate gills can evolve through repeated modification of a conserved endodermal tissue.

## Introduction

Vertebrate gills function either externally on the body surface or internally within the pharyngeal cavity, representing two distinct anatomical solutions for branchial respiration. Whether these contrasting respiratory organs share a common developmental basis remains unclear. Amphibians are uniquely suited to investigate this possibility because they transiently develop both external and internal gills during ontogeny (1). While the external gills support early larval respiration (2,3), they are retained throughout life in some salamanders (4) or become replaced by internal gills during frog development (5,6), providing an exceptional opportunity to investigate the developmental relationship between these two gill types.

External and internal gills in vertebrates differ markedly in their anatomy and developmental context (7,8). External gills arise as epithelial outgrowths from the hyoid or branchial arches, whereas internal gills develop within the pharyngeal apparatus as serial respiratory filaments. Outside amphibians, external gills are found only in lepidosirenid lungfishes and bichirs (9,10), where they occupy different arch positions and are generally considered to have evolved independently (11,12). In contrast, internal gills are widespread among aquatic vertebrates and are regarded as homologous structures derived from the ancestral vertebrate pharyngeal apparatus (13,14). Despite these anatomical and evolutionary differences, external and internal gills share remarkable structural and functional similarities (12,15–19), raising the question of whether they also share aspects of their developmental organization.

The embryonic origin of the epithelia forming vertebrate gills has long been central to this question. Fate-mapping studies in cartilaginous and ray-finned fishes have demonstrated that internal gills arise from pharyngeal endoderm (13,20–22), establishing this tissue as a conserved source of the vertebrate respiratory epithelium. By contrast, amphibian external gills have traditionally been regarded as purely ectodermal structures (18,23–28), despite classical embryological experiments showing that their development depends on the presence of the pharyngeal endoderm (29–34). Whether the pharyngeal endoderm merely induces external gill formation or forms a continuous epithelial component of the developing gills has remained unresolved. Resolving this question is essential for understanding the developmental relationship between external and internal gills.

Here, we combine histological analysis with direct lineage tracing in representatives of two major amphibian clades, the Mexican axolotl (*Ambystoma mexicanum*) and the African clawed frog (*Xenopus laevis*), to resolve the embryonic origin of amphibian gills. We show that the pharyngeal endoderm forms a continuous epithelial component of both external and internal gills, identifying a common epithelial framework underlying amphibian gill morphogenesis. These findings provide a developmental basis for understanding how diverse gill types can arise through repeated modification of a conserved endodermal tissue.

## Results

### Histological analysis suggests progressive expansion of the pharyngeal endoderm into the developing external gill

To determine the epithelial organization of the developing external gill, we first examined gill morphogenesis histologically in the axolotl. At the onset of external gill development, the eosinophilic, yolk-rich endoderm forms the epithelial lining of the pharyngeal wall, covering only the medial surface of the branchial arches (**Fig. 1A**). As the external gills begin to emerge, however, the endoderm progressively expands around the branchial arch mesenchyme, extending onto its lateral surface (**Fig. 1E, arrows**) and subsequently into the growing gill bud (**Fig. 1B, F, arrows**). Throughout gill outgrowth, the endoderm remains confined to the basal epithelial layer, while the apical layer is formed by ectoderm. As development proceeds, the endoderm extends distally to the tip of the growing external gill, with basal ectoderm persisting only in a narrow region immediately posterior to the tip and within the forming gill slits **(Fig. 1A, B, and E, F, light blue**). This epithelial organization is maintained throughout gill morphogenesis, including after elongation of the gill stem and formation of the gill filaments (**Fig. 1C, D, G, H**). Together, these observations suggest that the pharyngeal endoderm progressively expands into the developing external gill to form most of its basal epithelium, while the ectoderm consistently remains as the superficial epithelial layer.

**Figure 1.**
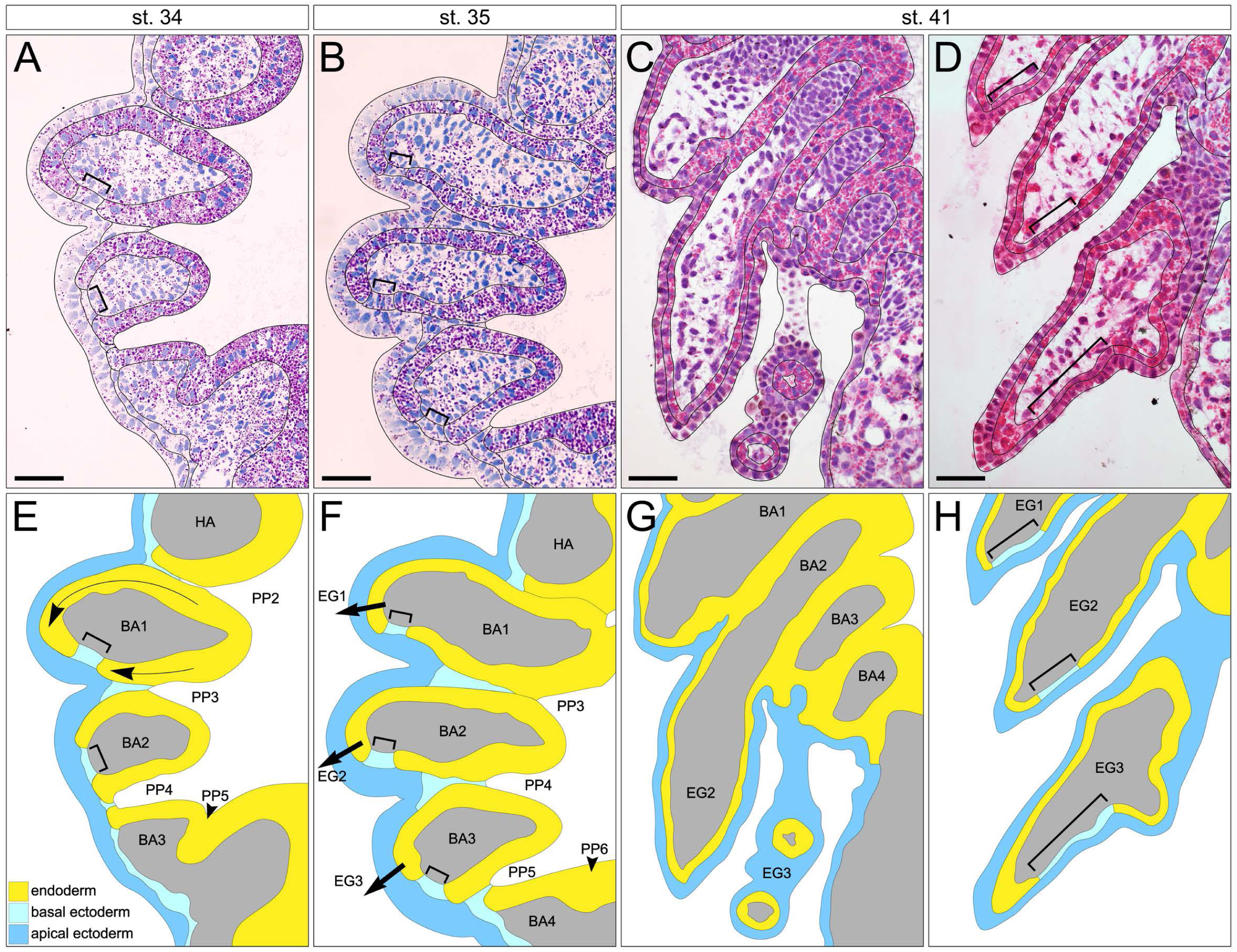
Histological analysis of external gill development in the Mexican axolotl. Horizontal sections through developing external gills stained with hematoxylin and eosin (A-D) and corresponding interpretations of ectodermal and endodermal distributions (E-H). During the initial stages of external gill outgrowth, eosin intensely stains the yolk-rich pharyngeal endoderm, allowing it to be distinguished from surrounding tissues (A, B). As development proceeds, this distinction gradually diminishes with yolk consumption, although endodermal cells remain identifiable (C, D). The interpreted sections illustrate progressive distal extension of the endodermal epithelium (yellow) along the branchial arches toward the developing external gills (E, arrows). During this process, the endoderm forms a continuous basal epithelial layer extending to the distal gill tips (F–H, arrows), whereas the ectoderm remains as the superficial epithelial layer (dark blue). Small regions of basal ectoderm persist immediately posterior to the gill tips (E, F, H, brackets) and at the sites of future pharyngeal slits. The mature external gill epithelium is therefore interpreted as a double-layered epithelium composed of an apical ectodermal layer and a basal endodermal layer, except in the small ectoderm-only regions posterior to the gill tips. Note that C and D are serial sections of the same specimen. Thin arrows in E indicate distal extension of the endodermal epithelium; arrows in F mark the direction of external gill outgrowth; brackets indicate regions of basal epithelium composed of ectoderm. BA, branchial arch; EG, external gill; HA, hyoid arch; PP, pharyngeal pouch. Scale bar = 100 µm.

### Lineage tracing demonstrates continuous endodermal contribution to the external gill epithelium

To directly test the histological observations, we traced the fate of the pharyngeal endoderm by injecting CDCFDA into the neurula-stage foregut cavity, thereby specifically labelling the foregut endoderm without contaminating surrounding mesoderm or ectoderm except at the injection site (**Fig. 2A**). In whole mounts, labelled cells were readily detected within the pharyngeal slits and extended into the developing external gills (**Fig. 2B**). As gill outgrowth progressed, the labelled endoderm formed a continuous epithelial domain that initially occupied the anterior portion of the gill bud (**Fig. 2C, arrow**), subsequently expanded towards the distal tip (**Fig. 2D, E, arrows**), and later also extended separately into the posterior portion of the gill (**Fig. 2D, E, arrowheads**), while consistently leaving a narrow unlabeled region immediately behind the gill tip (**Fig. 2F, G**). Histological sections confirmed that the labelled pharyngeal endoderm progressively expanded from the pharyngeal lining into the gill anlage from both its anterior and posterior aspects (**Fig. 2H–J, arrows and arrowheads**), ultimately forming most of the basal epithelial layer of both the gill stem and the gill filaments, whereas the superficial epithelial layer remained ectodermal (**Fig. 2J, K**). Notably, the absence of comparable epithelial expansion in the mandibular and hyoid arches suggests that continuous endodermal extension is a morphogenetic feature specific to branchial arches undergoing external gill formation.

**Figure 2.**
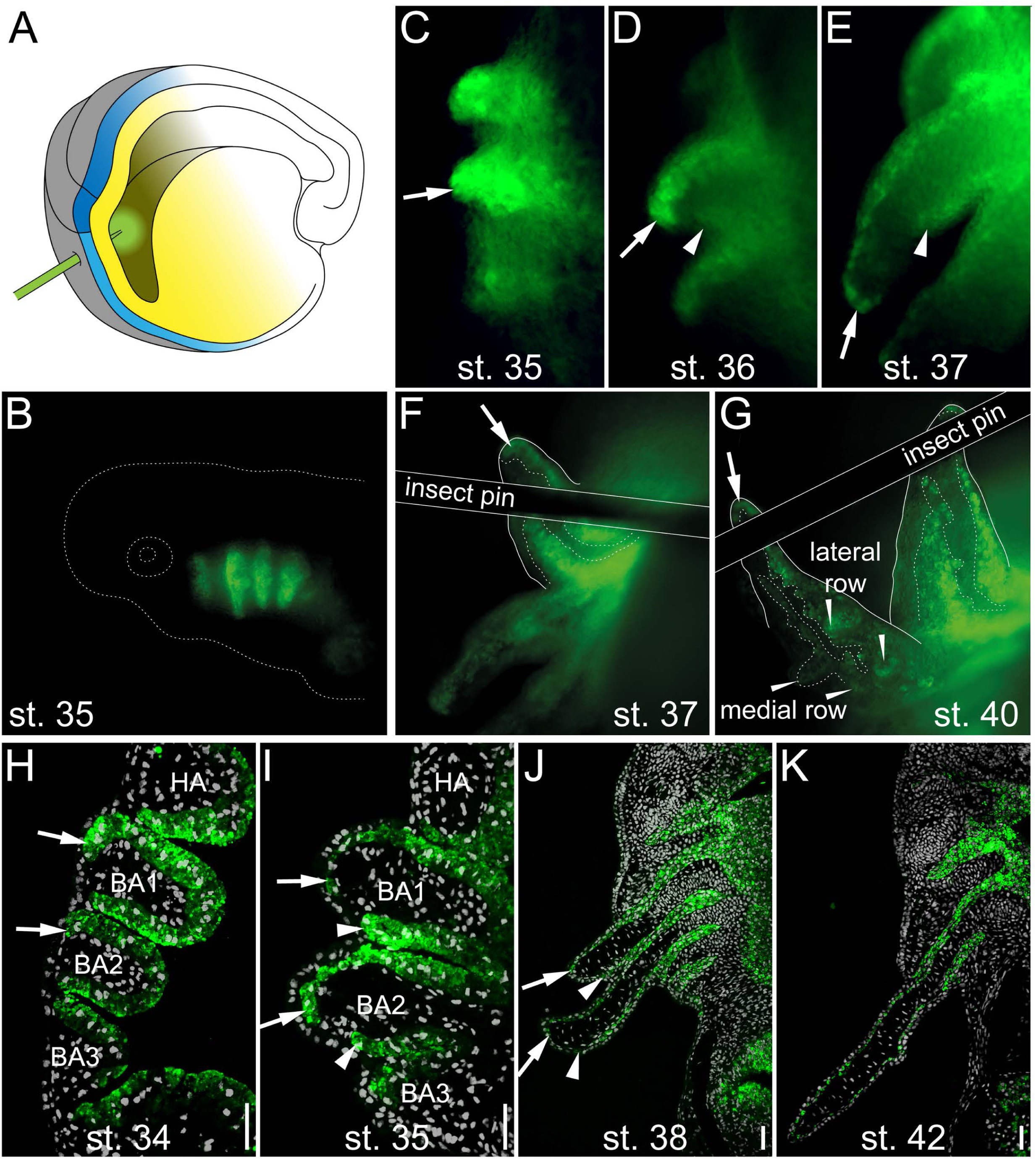
Fate mapping of pharyngeal endoderm during external gill development in the Mexican axolotl. Pharyngeal endoderm was labelled by injecting CDCFDA into the foregut cavity through the prospective mouth region of an early neurula embryo (A). Whole-mount preparations confirm specific labelling of the pharyngeal endoderm (B). During external gill outgrowth, labelled endoderm forms a continuous epithelial sheet extending along the anterior surface of the developing gill to its distal tip (C–E, arrows). A shorter endodermal extension subsequently develops along the posterior surface of the gill (D, E, arrowheads). When flipped, the posterior aspect of the external gill contains a characteristic Λ-shaped endoderm-free region immediately behind the gill tip (F, G). Horizontal sections demonstrate that the labelled endoderm extends distally from the pharyngeal epithelium along the anterior side of the branchial arch into the developing external gill (H, I, arrows), followed by a similar extension along its posterior side (I, J, arrowheads). The anterior and posterior epithelial sheets remain separated (J), leaving an endoderm-free region immediately posterior to the distal gill tip. Consequently, the basal epithelial layer of the external gill is formed predominantly by pharyngeal endoderm, whereas the superficial epithelial layer remains ectodermal (H–K). Arrows indicate the distal front of the anterior endodermal epithelium; arrowheads indicate the distal front of the posterior endodermal epithelium; thin arrowheads in G mark gill filaments; brackets indicate regions of basal epithelium composed of ectoderm. BA, branchial arch; HA, hyoid arch. Scale bars = 100 μm.

### Xenopus external gills exhibit the same epithelial organization as axolotl

To determine whether the epithelial organization identified in axolotl is conserved across amphibians, we examined external gill development in *Xenopus laevis*. Because the development of *Xenopus* external gills has remained poorly characterized, we first reconstructed their morphogenesis using µCT imaging combined with histological analysis. At NF stages 37–38, the three external gills appear as lateral protrusions of the branchial arches (**Fig. 3A**), and virtual sections reveal yolk-rich pharyngeal endoderm extending continuously from the pharyngeal cavity to the distal gill tip (**Fig. 3E**). Histological sections confirmed that these yolk-rich endodermal cells surround the branchial arch mesenchyme and occupy the growing gill epithelium (**Fig. 3I**). During subsequent gill elongation (**Fig. 3B**), the endodermal continuity is maintained, with yolk-rich cells remaining evident at the distal gill tip (**Fig. 3F, J**). As development proceeds, the external gills become enclosed by the opercular flap (**Fig. 3D**), while the branchial arches simultaneously differentiate into anterior filter rows and posterior internal gills (**Fig. 3G, H**). At these later stages, progressive depletion of yolk platelets prevents reliable histological discrimination of the pharyngeal endoderm from the surrounding epithelia (**Fig. 3K, L**). We therefore used direct lineage tracing to determine whether the endoderm remains continuous with both the transient external gills and the emerging internal gills.

**Figure 3.**
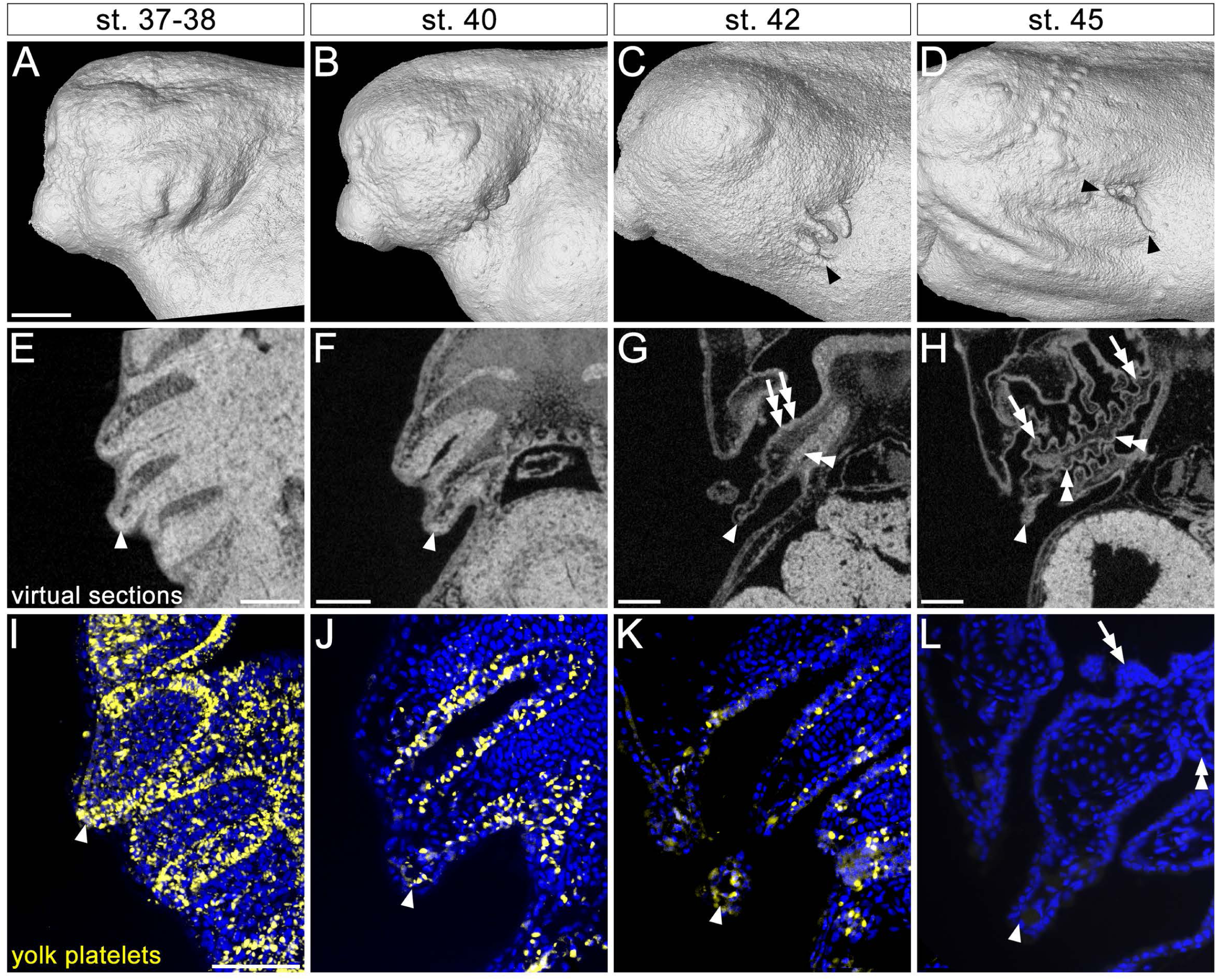
Morphological analysis of gill development in Xenopus laevis. MicroCT reconstructions illustrate the initiation and elongation of the external gills (A–C) and their subsequent enclosure by the opercular flap (C, D, black arrowheads). Virtual sections through the developing gills (E–H) reveal high contrast yolk-rich pharyngeal endoderm extending from the pharyngeal cavity to the distal gill tips during early stages (E, F). As the opercular flap forms, the branchial arches differentiate into anterior filter rows and posterior internal gills (G, H, double arrows and double arrowheads), while the yolk-rich endoderm gradually becomes indistinguishable owing to yolk consumption. Histological sections confirm the presence of autofluorescent yolk platelets within pharyngeal endoderm cells (I–L). At early stages, these cells line the pharyngeal cavity and extend to the tips of the external gills (I, J). As development proceeds, the yolk platelets are progressively depleted (K, L), preventing further identification of the endoderm based on morphology alone. Arrowheads indicate external gill tips; double arrows indicate filter rows; double arrowheads indicate internal gills. Scale bar in A = 200 μm (A–D); scale bars in E–H = 100 μm; scale bar in I = 100 μm (I–L).

### Lineage tracing demonstrates continuous endodermal contribution to both external and internal gills

To determine whether the pharyngeal endoderm remains continuous with both the transient external gills and the subsequently forming internal gills, we performed foregut lineage tracing in *Xenopus laevis* using CDCFDA labelling (see **Fig. 2A**). Labelled pharyngeal endoderm was readily detected in the pharyngeal slits and extended into the developing external gills (**Fig. 4A, B**). Viewed dorsally, labelled cells occupied the posterior surface of the external gills and reached their distal tips (**Fig. 4C, arrowheads**), providing whole-mount evidence that the continuous endodermal epithelium extends throughout the outgrowing gill. Horizontal sections confirmed that the labelled endoderm forms a continuous epithelial sheet extending laterally from the pharyngeal lining through the pharyngeal pouches into the posterior epithelium of the external gills, reaching their distal tips (**Fig. 4D–F**). As the branchial apparatus differentiated, the labelled endoderm remained continuous with the epithelia of both the anterior filter rows (**Fig. 4G**) and the posterior internal gills (**Fig. 4H**). Together, these observations identify the frog external gill as a direct epithelial extension of the pharyngeal endoderm that remains continuous with the epithelium giving rise to the internal gills.

**Figure 4.**
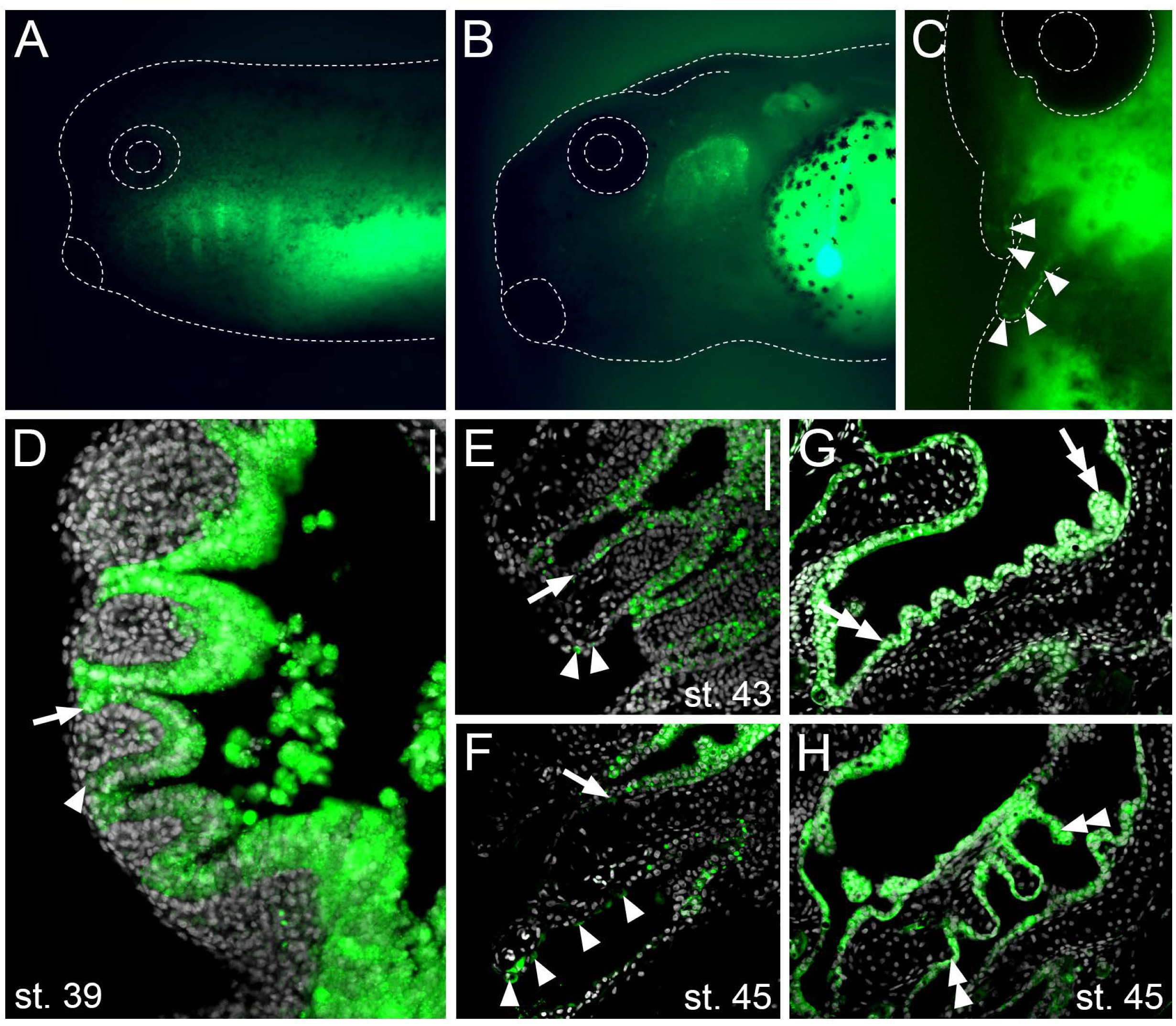
***Fate mapping of pharyngeal endoderm during gill development in* Xenopus laevis.** Pharyngeal endoderm was labelled by injection of CDCFDA into the foregut cavity as shown in Figure 2A. Whole-mount preparations confirm specific labelling of the pharyngeal endoderm, visible externally as a series of fluorescent domains at sites where the pharyngeal endoderm contacts the ectoderm (A). At later stages, labelled endoderm extends into the developing external gills (B). A dorsal view shows that the labelled cells populate the posterior epithelium of the external gills and reach their distal tips (C). Horizontal sections demonstrate continuous labelling of the pharyngeal endoderm surrounding the branchial arches and contacting the ectoderm at the prospective gill slits (D). The labelled endoderm extends into the posterior epithelium of the external gills, reaching their distal tips (E, F, arrowheads), whereas the anterior epithelial surface remains unlabelled (E, F, arrows). During differentiation of the pharyngeal apparatus, the labelled endoderm forms the epithelial lining of both the anterior filter rows (G, double arrows) and the posterior internal gills (H, double arrowheads), demonstrating epithelial continuity between the transient external gills and the definitive internal gills. Note that G and H are serial sections of the same specimen. Arrows indicate the distal front of the anterior endodermal epithelium; arrowheads indicate the distal front of the posterior endodermal epithelium; double arrows indicate filter rows; double arrowheads indicate internal gills. Scale bar in D = 100 μm; scale bar in E = 100 μm (E–H).

## Discussion

Here, we show that amphibian external gills develop as a continuous epithelial extension of the pharyngeal endoderm rather than as purely ectodermal outgrowths. This finding resolves a longstanding controversy over the embryonic origin of external gills and provides a developmental explanation for classical embryological experiments demonstrating that removal of the pharyngeal endoderm abolishes external gill formation (29–34). Because the pharyngeal endoderm contributes directly to the developing gill epithelium, these experiments can no longer be interpreted solely as evidence for an inductive role of the endoderm but also reflect the loss of a structural component of the developing organ. Whether the pharyngeal endoderm additionally provides molecular signals that regulate external gill outgrowth remains an important question for future studies.

The continuous contribution of pharyngeal endoderm to the external gill provides a developmental link between external and internal gills. Although these structures differ in their position relative to the pharyngeal cavity and have traditionally been considered distinct morphogenetic entities (23), our results demonstrate that they share a common epithelial foundation. In *Xenopus*, the same endodermal epithelium that extends into the transient external gills subsequently builds the filter rows and internal gills that develop within the pharyngeal apparatus. This developmental continuity is consistent with previous fate-mapping studies in cartilaginous and ray-finned fishes showing that internal gills arise from pharyngeal endoderm (13,20–22), supporting a conserved role of pharyngeal endoderm in vertebrate gill formation. Interestingly, while the spatial progression of endodermal extension differs between axolotl and *Xenopus*, both species ultimately establish the same epithelial organization, with the endoderm reaching the distal region of the external gill while the ectoderm remains superficial. Thus, species-specific morphogenetic trajectories appear to modify the implementation of external gill development without altering the underlying epithelial relationship.

The conserved epithelial relationship identified in amphibians provides a new perspective on the evolution of external gills across vertebrates. External gills are generally considered to have evolved independently in amphibians, lungfishes, and bichirs due to their distinct anatomical positions and the absence of clear fossil evidence linking these structures (11,12). However, despite their supposed independent evolutionary histories, external gills in these lineages appear to share a common developmental feature: the involvement of pharyngeal endoderm in their formation. Previous observations have suggested endodermal participation in the development of lungfish and bichir external gills (10,35), consistent with the possibility that pharyngeal endoderm represents a conserved developmental resource repeatedly recruited during external gill formation. Rather than requiring the evolution of a novel epithelial source, the emergence of external gills may therefore have relied on modification of an existing pharyngeal endodermal program capable of generating respiratory epithelia in diverse anatomical contexts.

The repeated involvement of pharyngeal endoderm in the formation of different gill types highlights a broader developmental property of vertebrate respiratory organs. Rather than representing unrelated epithelial structures assembled from distinct embryonic sources, internal and external gills appear to arise through modification of a shared endodermal developmental capacity. This perspective is consistent with the endodermal origin of other major vertebrate respiratory organs, including the lungs, which develop from foregut endoderm. Together, these findings suggest that pharyngeal and foregut endoderm provide a conserved epithelial foundation that can be repeatedly remodeled to generate respiratory structures adapted to diverse anatomical and ecological contexts. By revealing continuous endodermal participation in both external and internal gills, our study provides a developmental framework for understanding how vertebrate respiratory organs diversify while retaining a common embryonic origin.

## Materials and methods

### Embryo handling

Axolotl (*Ambystoma mexicanum*) embryos were obtained from the colony maintained at the Department of Zoology, Charles University, Prague, Czech Republic. *Xenopus laevis* embryos were obtained from colonies maintained at the Institute of Biotechnology, Czech Academy of Sciences, Vestec, Czech Republic, and the Department of Experimental Biology, Faculty of Science, Masaryk University, Brno, Czech Republic. Animal husbandry was performed in accordance with EU Directive 2010/63/EU on the protection of animals used for scientific purposes and Czech Act No. 246/1992 on the protection of animals against cruelty. All experiments were conducted in accordance with Czech legislation governing the use of animals in research and were approved by the relevant institutional and governmental authorities (MSMT-30784/2022 and MSMT-21426/2025, Ministry of Education, Youth and Sports of the Czech Republic; 45055/2020-MZE-18134, 45980/2023-MZE-13143, 31255/2019-MZE-18134, and MZE-20788/2025-13143, Ministry of Agriculture of the Czech Republic; MZP/2025/630/2482, Ministry of the Environment of the Czech Republic).

Axolotl embryos were obtained by natural spawning as previously described (36). After completion of spawning, embryos were collected, rinsed in tap water, maintained in sterile 1× Steinberg solution supplemented with antibiotics, and manually decapsulated. *Xenopus* embryos were generated and maintained according to established protocols (37). Briefly, adult males were anesthetized in 20% MS-222 (Sigma-Aldrich, A5040), and testes were surgically excised and transferred to ice-cold 1× Marc’s Modified Ringer’s solution (MMR) supplemented with 50 µg/mL gentamicin (Sigma-Aldrich, G3632). Ovulation in sexually mature females was induced by injection of 260 U human chorionic gonadotropin (Sigma-Aldrich, CG5) into the dorsal lymph sac approximately 16 h prior to egg collection. Females were maintained overnight at 18 °C. Eggs were collected by gentle abdominal pressure and fertilized *in vitro* using freshly macerated testis tissue in 0.1× MMR. Jelly coats were removed using 2% cysteine, and embryos were maintained in 0.1× MMR supplemented with antibiotics.

### Dye administration and endoderm fate mapping

Stage 14 axolotl and Nieuwkoop–Faber (NF) stage 13 *Xenopus* embryos (38,39) were positioned in cavities made in modelling clay and subjected to dye injection. Pharyngeal endoderm was fate-mapped using CDCFDA carboxyfluorescein (AAT Bioquest), a fluorescent cell-labeling compound previously used to label epithelial cell sheets (40). A 50 mM stock solution of CDCFDA was diluted 1:10 in 10% sucrose. The resulting solution was injected into the foregut cavity of neurula-stage embryos. The injection site was positioned below the transverse neural fold, allowing the capillary to enter through the prospective mouth region. This approach was designed to label prospective pharyngeal endoderm while avoiding mesoderm and ectoderm, apart from a small number of oral ectodermal cells at the injection site. Following injection, embryos were transferred individually to fresh medium in agarose-covered 24-well plates and screened for green fluorescence. At selected developmental stages, embryos were anesthetized by overdose with MS-222 (Sigma-Aldrich) and fixed overnight in 4% paraformaldehyde (PFA) in 0.1 M phosphate-buffered saline (PBS) for subsequent analysis (axolotl, n = 14; *Xenopus*, n = 32).

### Plastic histology and imaging

For histological analysis, embryos were embedded in JB-4 plastic resin (Polysciences) according to the manufacturer’s instructions and sectioned horizontally at 10 µm thickness. Sections from untreated embryos were stained with hematoxylin and eosin, mounted in DPX mounting medium (Sigma-Aldrich), and coverslipped. Sections from CDCFDA-injected embryos were mounted in Fluoroshield mounting medium containing DAPI (Sigma-Aldrich). Images were acquired using an Olympus BX51 microscope equipped with a DP74 camera and CellSens software. Whole-mount CDCFDA-injected embryos were imaged using a Zeiss SteREO Lumar.V12 stereomicroscope.

### Micro-computed tomography scanning

*Xenopus* embryos at NF stages 37–38, 40, 42, and 45 were fixed in 4% PFA in 0.1 M PBS. To enhance soft-tissue contrast, samples were stained in Lugol’s solution diluted 1:1 with distilled water for 24 h. Samples were immobilized for scanning by embedding them in 1% agarose within 1.5-mm glass capillaries. Micro-computed tomography (µCT) scans were acquired using a Phoenix Nanotom M laboratory µCT system with the following parameters: 90 kV, 150 µA, 1000 ms exposure time, 2100 projections, 35 min scan time, 0.1-mm Al filter, and 0.8-µm voxel resolution. Data were reconstructed using phoenix datos|x 2.0 software, and all data visualization was performed using VGStudio MAX 2025.3.

## Acknowledgments

We thank Adéla Dienstbierová and Anna Hirnerová for axolotl colony care, and Douglas Porto for *Xenopus* colony care. This work was supported by the Czech Science Foundation (GACR 23-07212S) to HS and VS, Grant Agency of the Charles University (GAUK 179524 and 594561) to HS and JV, and Faculty of Science Charles University STARS PhD program to HS. RŠ was supported by RVO: 86652036. We acknowledge the CREATIC project funded by the European Union (Grant Agreement No. 101059788).

## Author contributions

Conceptualization: VS

Data curation: HS, JV, MK, VS

Formal analysis: MK

Funding acquisition: HS, JV, VS

Project administration: VS

Visualization: HS, MK, JV, VS

Methodology: HS, JV, MK, VS

Resources: LAM, JH, JK, RŠ, VS

Supervision: JH, JK, RŠ, VS

Writing - original draft: VS

Writing - review and editing: LAM, JH, RŠ, VS

All authors read and approved the final version of the manuscript.

